# Asymmetric mitochondrial trafficking maintains network morphology by balancing perinuclear biogenesis

**DOI:** 10.64898/2025.12.08.692722

**Authors:** Julius Winter, Juan C. Landoni, Keaton Holt, Sheda Ben Nejma, Emine Berna Durmus, Tatjana Kleele, Elena F. Koslover, Suliana Manley

**Author notes:** These authors contributed equally: Julius Winter, Juan C. Landoni.

## Abstract

In functionally polarized cells, mitochondria can form distinct subpopulations, positioned at sites of varying metabolic and energetic demands. Thus far, the potential presence of such subpopulations and implications of their intracellular trafficking in immobile and proliferative cells remains largely undescribed, despite such cells serving as key models. Here, we use substrate micropatterning to create reproducible morphologies of cultured immortalized cells, enabling us to define mitochondrial subpopulations and follow their trafficking by photoactivation. We discovered that mitochondrial material is dispersed asymmetrically throughout the cell via biased anterograde transport from the perinuclear area. Combining quantitative analysis and *in silico* modeling, we characterize the causes and consequences of unbalanced mitochondrial trafficking. Our findings indicate that this bias is required to distribute new material resulting from perinuclear mitochondrial biosynthesis to sustain mitochondrial mass distribution across the cell, and to maintain normal network connectivity.

## Introduction

Most mammalian cells contain tens to hundreds of individual mitochondria, which vary in size and connectivity (Collins et al., 2002; Landoni et al., 2024). This morphological plasticity, when coupled with organelle transport and anchoring, allows mitochondria to distribute their function throughout the cell (Fawcett, 1981; Mironov, 2007). Mitochondria are frequently positioned at sites of specific signaling, energetic, or metabolic demand, including the leading edge of migrating cancer cells (Cunniff et al., 2016), the immunological synapse of T-cells (Quintana and Hoth, 2012; Quintana et al., 2007), contacts with lipid droplets (Valm et al., 2017) or intracellular parasites (Matsumoto et al., 1991; Medeiros et al., 2025), and axonal synapses (Rangaraju et al., 2014).

Axonal mitochondrial trafficking occurs over distances in the range of meters, making it one of the most extreme cases of transport in cell biology, whose impairment can underlie human neurodegenerative diseases (Sheng and Cai, 2012). Consequently, longrange mitochondrial transport has been well-characterized in neurons, where it is mediated by microtubule motors (Misgeld et al., 2007). The outer mitochondrial membrane proteins MIRO1 and MIRO2 (Morlino et al., 2014; Wang and Schwarz, 2009) associate mitochondria to motor proteins in a calciumdependent manner via the adaptor proteins TRAK1 (tethering to kinesins), and TRAK2 (tethering both kinesin and dynein) (Fenton et al., 2021; Glater et al., 2006; van Spronsen et al., 2013). While mitochondria traffic bidirectionally along axonal microtubules (Misgeld and Schwarz, 2017; Saxton and Hollenbeck, 2012), anterograde-moving organelles present a higher membrane potential than their retrograde-moving counterparts, highlighting the role trafficking plays in supplying healthy organelles throughout the cell (Lin et al., 2017; Miller and Sheetz, 2004; Zheng et al., 2019).

Many cell types lack such extreme polarized function and architecture. Yet, to what extent mitochondrial active transport and life cycle processes (mitochondrial dynamics and DNA replication) are spatially organized in compact cells lacking polarized function remains largely unknown. Here, we address this knowledge gap in immobile fibroblast-like cells (COS-7), which are typically unpolarized. By culturing the cells on patterned coverslips, we established a platform for reproducible quantification of mitochondrial transport across cells, leading to the discovery that content mixing occurs asymmetrically from the perinuclear region towards the periphery. Combined with *in silico* modelling, we gain insights into the interplay between biased transport and local biogenesis and predict their impacts on mass distribution, as well as network renewal and morphology. We perturbed relative transport rates by modulating motor protein adaptors TRAK1 and TRAK2, and found changes in mitochondrial distribution and connectivity consistent with our predictions. Our findings emphasize the role of active transport in maintaining mitochondrial homeostasis, not only by distributing new material, but also by sculpting organelle connectivity.

## Results & Discussion

### Anterograde transport of small mitochondria is dominant in COS-7 cells

Fibroblast-like COS-7 cells are morphologically heterogeneous and compact (Fig. 1A), and when immobile, lack apparent polarized function. Thus, intracellular transport in these cells is challenging to quantify, in contrast to the simplified analysis of one-dimensional organelle motion in neuronal axons. To obtain consistent measurements of mitochondrial transport, we cultured COS-7 cells on micropatterned coverslips, resulting in relatively uniform shapes (Fig. 1B-C) with a well-defined cytoskeletal architecture and a shared reference frame (Chevrollier et al., 2012). Patterning also immobilized cells, maintaining their morphology and position over time, enabling long-term measurements of equivalent subcellular regions. After screening a variety of shapes, we chose the arch, or “empanada”, which consistently yielded a good number of cells (~25% of patterns occupied) with reproducible morphologies consisting of a crescent with two tips (Fig. 1B-C). The resulting cell boundary has an axis of symmetry with the nucleus, near the center axis, shifted towards one of the cell tips. Mitochondrial density is enriched in the perinuclear region towards the other cell tip, likely associated with the microtubule-organizing center, and henceforth referred to as the perinuclear cluster (PNC).

**Figure 1:**
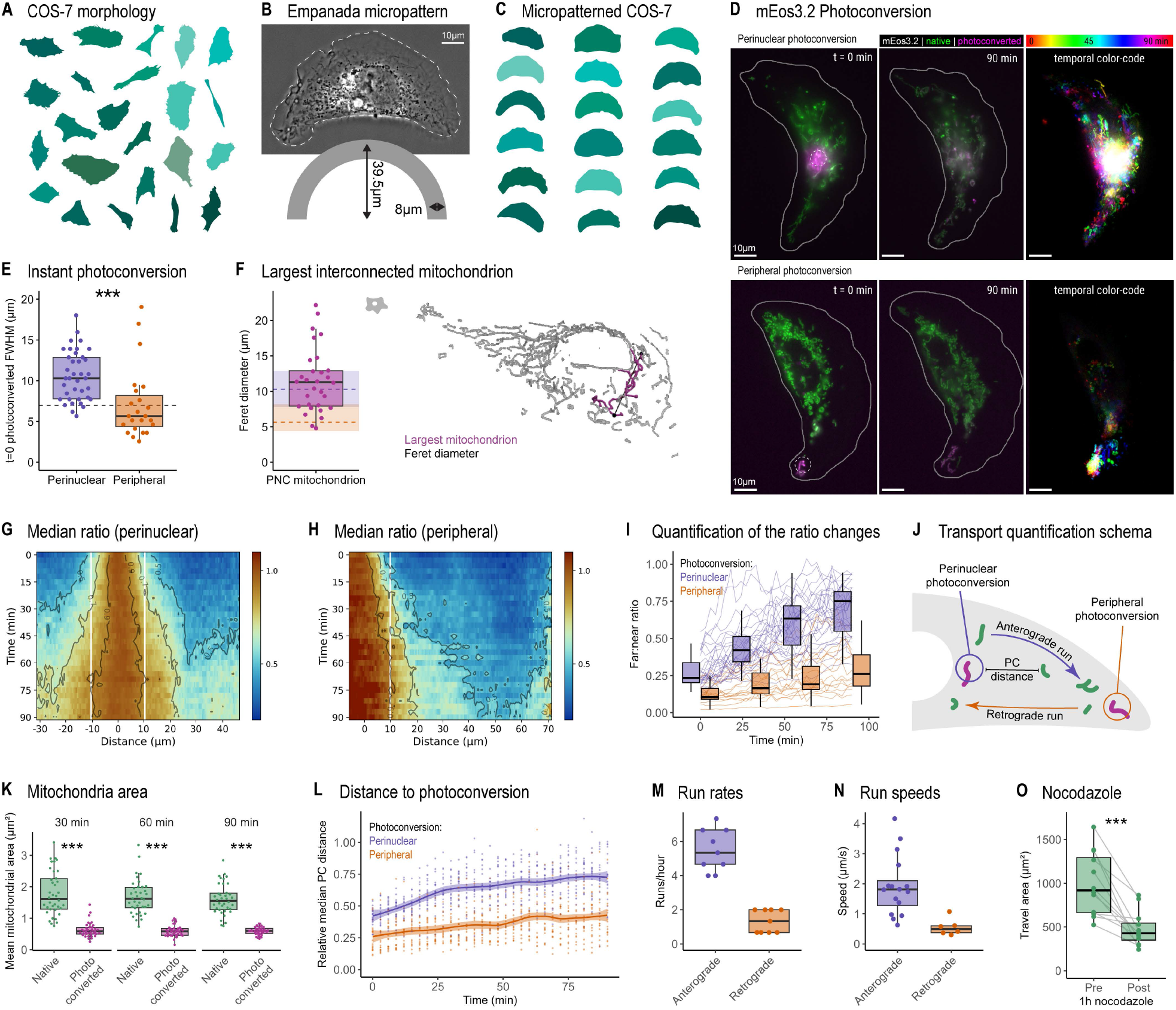
Anterograde transport of small mitochondria is dominant in COS7 cells. (A) Montage of COS-7 cell shapes from plain glass coverslips. (B) Representative phase-contrast image of a patterned cell (top), and geometry of the arch-shaped “empanada” micropattern, used for cellular patterning (bottom). (C) Montage of COS-7 cell shapes from micropatterned coverslips. (D) Representative cells from perinuclear and peripheral photoconversion experiments. Merged native and photoconverted mEos3.2 (green and magenta, respectively) from first and last frames, and temporal color-code of the photoconverted channel across the entire time series (right). Dotted circle, approximate photoconversion region. (E) FWHM of instantaneous photoconverted regions (t=0). Dashed line indicates the theoretical size of photoconversion beam. (F) Feret’s diameter of the largest mitochondrial cluster from volumetric segmentation (as in the right). Shaded dashed lines indicate median and inter-quartile ranges of photoconversion FWHM as in (E), ameboid symbol indicates the data comes from non-empanada micropatterned cells. (G-H) Median of ratios kymograph for perinuclear (G, N=29 cells) or peripheral (H, N=16 cells) photoconversion. Horizontal axis spans the tip-to-tip cell extent, zero marks the site of photoconversion, vertical white lines indicate the 11 μm instantaneous diffusion boundary. (I) Ratio between the median values beyond (far) and within (near) the instantaneous diffusion boundary as in (G-H). (J) Schema of quantified measures. (K) Comparison between the area of native and photoconverted mitochondria outside the photoconversion area at select time points; points represent the median of individual cells (N=38 cells). (L) Median distance of single photoconverted mitochondria to the photoconversion zone, normalized to the median distance of native mitochondria. Each data point represents the relative median for an individual cell (N=38 cells). Smoothed conditional means and 95% confidence interval for peripheral and perinuclear photoconversion experiments as shaded lines. (M) Directed mitochondrial runs in either direction; each point represents a single cell’s rate in runs/h (N=9 cells). (N) Single mitochondria velocities extracted from high-speed images and manual tracking (N=16+6 anterograde vs retrograde mitochondria from two independent cells). (O) Mitochondrial movement with and without microtubule integrity, calculated as segmented overlap between timeframes, before and after 1h nocodazole incubation. P values from two-sided Wilcoxon rank sum test (p< 0.05*, 0.01**, 0.001***).

To visualize transport dynamics and content mixing, we expressed the mitochondrial matrix-targeted photoconvertible (green-to-red) fluorescent protein mEos3.2 (mito-mEos3.2) in micropatterned cells. A small circular region (area *A* = 38.5 µm^2^, radius *r* = 3.5 µm) of mitochondria was partially photoconverted (Fig. 1D), either within the PNC or at a cell tip (Fig. S1A), followed by time-lapse imaging (3 min/image). Photoconversion at the two locations presented remarkably different mixing behaviors: Peripheral mitochondria rarely underwent retrograde transport, and material mostly remained in a small volume near the cell edge (Fig. 1D). In contrast, perinuclear material mixed nearly instantaneously within the PNC network, followed by the emergence of small mitochondria which moved throughout the cell, fusing with other mitochondria including at the cell periphery within 90 min (Fig. 1D, S 1A-B, Movie S1). Taking advantage of the reproducible shape of the cells, we sought to quantify the spreading of material. For each cell and time point, we integrated the photoconverted intensity over the cell’s peripheral width at each location along the central arch to obtain a 1D intensity profile (Fig. S1C-D). As observed above, perinuclear photoconversion instantly spread beyond the circular photoconverted region (t=0) to an area significantly larger than that of peripheral photoconversion (Fig. 1E), implying rapid diffusion of proteins within the highly connected PNC mitochondria. The 10.3 ± 3.1 μm median diameter of the instantaneously photoconverted region also coincides with the 11.6 ± 4.5 μm median Feret’s diameter of the largest interconnected component in 3D mitochondrial network reconstructions (Fig. 1F, methods), further suggesting that the interconnected PNC network supports rapid diffusion.

To account for the underlying spatial variability in organelle abundance, we normalized the photoconverted intensity by the total signal (photoconverted + “native” (non-photoconverted)) to obtain a “ratio” value. We visualized the spread of material through time in ratio kymographs (Fig. S1E). Furthermore, we combined ratio kymographs across cellular replicates to obtain population medians (Fig. 1G-H). Constant ratio contours for each of the two photoconversion configurations revealed more extensive spreading of the photoconverted material towards the cellular edge in the perinuclear compared with the peripheral kymograph. We quantified this long-term spreading by using the calculated ± 11 μm from the photoconversion spot (white vertical lines, Fig. 1G-H) as a threshold or boundary for instantaneous diffusion through interconnected mitochondria, and measuring the relative median signal outside (far) vs. within (near) this area (Fig. 1I). The divergence of these ratios over ~90 minutes reflects that spreading from the perinuclear area is better homogenized throughout the cell compared with spreading from the cell periphery.

Fusion and network connectivity are known to play key roles in distributing mitochondrial material across the cell (Liu et al., 2009; Hoitzing et al., 2015). However, those processes result in unbiased, bilateral mixing, and therefore would not suffice to explain the asymmetry of material spreading we observe. We speculate that this difference arises from the small mitochondria that are rapidly transported from the perinuclear area, which lack a significant peripheral counterpart.

To quantify mitochondrial transport by an orthogonal method, we segmented individual mitochondria. We found that mitochondria moving out of the perinuclear photoconversion zone are smaller on average than the overall population (Fig. 1J-K), consistent with previous reports of higher motility in small mitochondria (Misgeld et al., 2007; Schwarz, 2013). We then measured the median of photoconverted mitochondria distances from the photoconversion region (normalized to the median distance for non-photoconverted mitochondria, Methods). This grows more rapidly with time for perinuclear than for peripheral photoconversion experiments (Fig. 1L), consistent with our kymograph analysis.

To further quantify these fluxes, we manually counted mitochondria presenting fast directed motion in either direction (anterograde movement towards the periphery, retrograde movement towards the perinuclear area). This resulted in an anterograde run rate of 0.091 ± 0.021 mitochondria/min and a retrograde rate of 0.022 ± 0.011 mitochondria/min (Fig. 1M). Using faster frame rate movies (2 s/image) following perinuclear photoconversion, we also measured the velocities of solitary motile mitochondria during brief bursts of directed motion. The velocities (anterograde = 1.81 µm s^-1^, retrograde = 0.50 µm s^-1^) (Fig. 1N, Movie S2) are consistent with values previously reported in neurons (0.5 - 3.5 µm s^-1^) and faster than mitochondrial displacement by actin comet tails (~0.2 µm s^-1^) (Moore et al., 2021), suggesting translocation by microtubule motor proteins. Indeed, the acute disruption of microtubule polymerization using nocodazole halts mitochondrial movement (Fig. 1O), detecting a median of 0 anterograde runs.

The empirical values of motile mitochondrial size and run rates allowed us to estimate the mitochondrial volume transferred from the PNC to the periphery (Methods), which resulted in a transport rate of ~1.2 μm^3^/h, or 48 μm^3^ of mitochondria per cell cycle. This transport capacity aligns closely with the calculated median peripheral mitochondrial volume of 47 μm^3^ (Methods), indicating that the measured anterograde trafficking is sufficient to account for the steady-state peripheral mitochondrial mass generation across the cell cycle.

Taken together, our data indicate that the spread of mitochondrial content in COS-7 cells is highly asymmetric. Peripheral mitochondria remain confined near the cell edge with limited retrograde trafficking activity, while perinuclear material mixes rapidly (< 1 sec) via diffusion throughout the interconnected PNC network, and spreads further toward the cell periphery over tens of minutes via the microtubular anterograde transport of small mitochondria.

### Mitochondrial biogenesis occurs preferentially in a perinuclear subpopulation

We wondered what function the asymmetric spreading of subcellular mitochondrial populations could serve. In neurons, mitochondrial biogenesis occurs preferentially in the cell body, and high-membrane potential organelles are actively trafficked to axons and dendrites via microtubule motor proteins (Ferree et al., 2013; Misgeld and Schwarz, 2017). A fraction returns via retrograde transport to complete the mitochondrial life cycle of biogenesis and quality control (Harbauer et al., 2022; Lin et al., 2017; Maday et al., 2012; Miller and Sheetz, 2004). Despite the lack of equivalent specialized cellular domains in non-polarized cells, we speculated that the observed asymmetric spreading could serve a similar purpose in the mitochondrial life cycle; thus, we investigated the spatial organization of mitochondrial biogenesis in COS-7 cells.

As mentioned, perinuclear mitochondria form an interconnected, networked cluster (PNC), whose size can be quantified using volumetric analysis (Fig. 1H & 2A). In contrast to this cluster, small mitochondria (< 2 µm^3^) are distributed throughout the cell (Fig. 2B), with those nearest to the edge of the cell exclusively within this small size range. Detecting and classifying spontaneous mitochondrial fission events (Methods) revealed an additional difference between these two cellular regions: while peripheral divisions occurred throughout the cell, midzone divisions were enriched within the PNC (Fig. 2C). We previously reported that midzone divisions are associated with mitochondrial proliferation (Kleele et al., 2021), which prompted us to test whether mitochondrial DNA (mtDNA) replication is also enriched at the perinuclear area.

**Figure 2:**
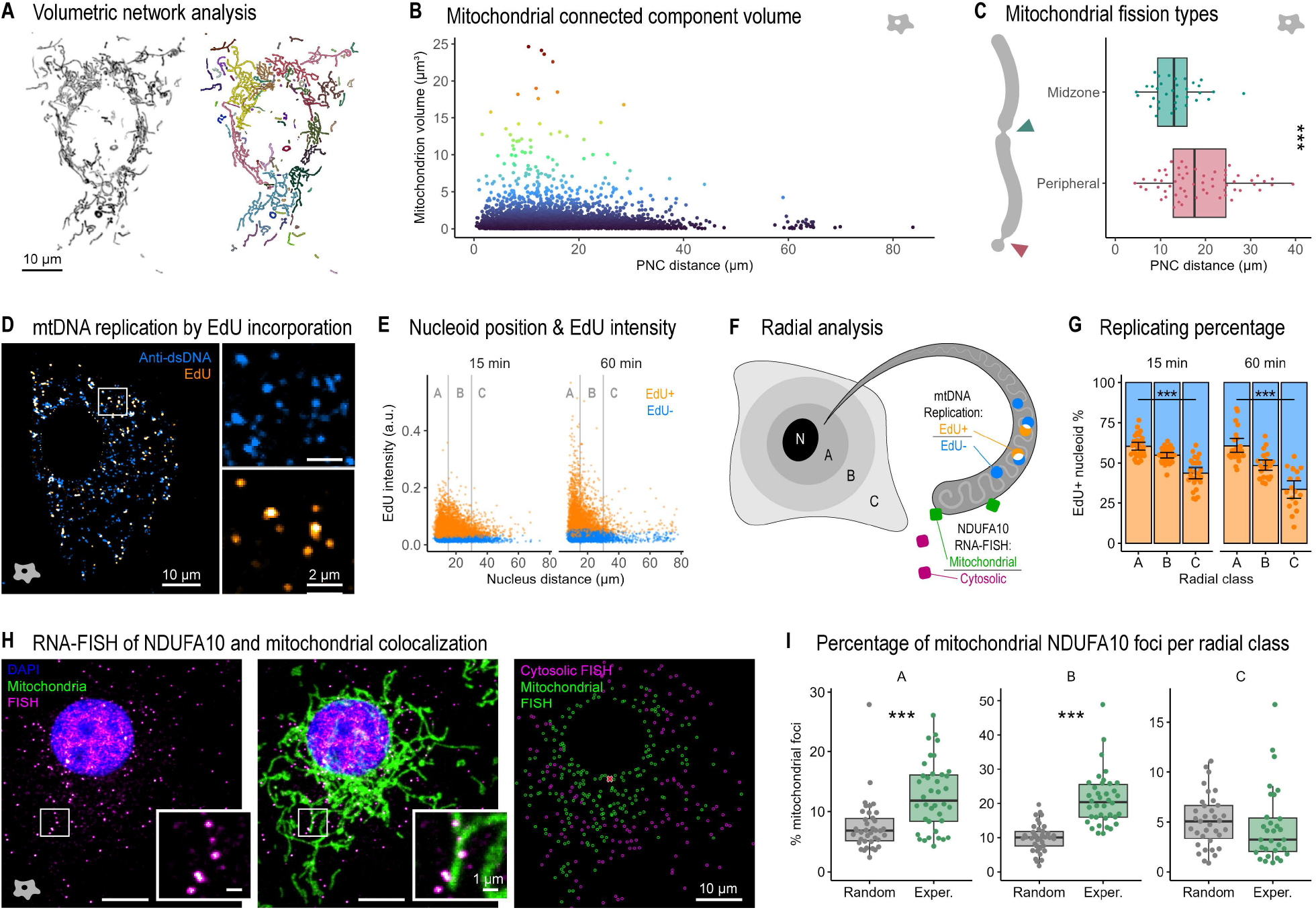
Mitochondrial proliferation is spatially segregated. (A) Representative mitochondrial network volume (mito-mScarlet, black) and segmentation, colored by connected components. (B) Volume of segmented mitochondria vs. distance from the PNC (N=19 cells, 5806 mitochondria). (C) Fission types and their distance to the PNC (peripheral position as <30% of the total mitochondrial length), distance and classification blinded of one another (N=10 cells, 94 fissions). (D) Representative nucleoid staining, anti-dsDNA antibody (blue) and click-detected EdU (orange) following a 60-min pulse, with separate channels as insets. Merge and insets include nucleus masking and top-hat filtering to enable nucleoid visualization (N_15min_=29 cells, N_60min_=22 cells). (E) Scatterplot of each detected nucleoid (from dsDNA immunostaining), distance to the nucleus vs. EdU intensity. Colors indicate their EdU positivity classification used for subsequent analyses. Vertical bars mark the radial analysis thresholds. (F) Graphical description of radial analysis and quantification approaches used subsequently; N = Nucleus. (G) Percentage of EdU+ nucleoids in each of the radial zones as in (D-E). (H) Representative NDUFA10 RNA-FISH (magenta), TOM20 immunostaining (green) and nucleus (blue). On the right, detected FISH foci classified based on their co-localization with mitochondria (N=38 cells, NFISH=11788 foci), and PNC marked in red. (I) Radial analysis of NDUFA10 RNA-FISH as in (F), percentage of mitochondrial foci per radial zone compared to an equivalent random positioning on identical networks. P values from two-sided Wilcoxon rank sum test (p< 0.05*, 0.01**, 0.001***), scale bar 10 µm. Ameboid symbol indicates the data comes from nonempanada-micropatterned cells.

We used the incorporation of the nucleotide analogue ethynyldeoxyuridine (EdU) to identify mtDNA nucleoids that underwent replication during 15- or 60-minute pulses. We labeled nucleoids using an anti-DNA antibody, imaged and segmented them, then analyzed the integrated intensity of EdU within those defined regions. We found an inverse relationship between EdU incorporation and distance from the nucleus (Fig. 2D-E). Radial analysis (Fig. 2F) further demonstrated that the perinuclear region is enriched in replicating nucleoids, while the cell periphery is depleted compared to the mean (Fig. 2G), with these differences present for both measured time windows.

Mitochondrial biogenesis also requires nuclear-encoded proteins that are translated in the cytosol, often carried out by ribosomes located near the mitochondrial surface for efficient import (Matsumoto et al., 2012). We determined sites of translation of the respiratory complex 1 subunit NDUFA10 by RNAFISH and found a sub-population that co-localized with mitochondria (Fig. 2H), consistent with previous reports (Matsumoto et al., 2012). To account for the increased density of mitochondria in the perinuclear area, we considered only foci that co-localized with mitochondria in both the experimental and random datasets. We performed radial analysis using the previously determined 11 μm PNC radius to define cellular regions, including the perinuclear region “A” and periphery “C”. This analysis revealed a significant enrichment of mitochondria-associated NDUFA10 mRNA foci in the perinuclear areas, ~2-fold higher than random, in contrast to the cell periphery (Fig. 2I).

Altogether, our data reveal the existence of two distinct mitochondrial subpopulations in unpolarized cells, with smaller organelles in the cell periphery, and an interconnected, perinuclear cluster exhibiting signatures of biogenesis and proliferation.

### Biased flow maintains network mass balance and connectivity *in silico*

While the outward flow of mitochondria could serve the purpose of spreading newly synthesized material, this was counterintuitive for a system we assumed to be at steady state. To gain mechanistic insight into the underlying dynamics, we modeled the mitochondrial network using a spatially resolved 3D graphbased formulation representing mitochondria as edges delimited by nodes (Holt et al., 2024). The simulation uses fission and fusion rates tuned to produce networks with experimentally determined connectivity parameters, inside a geometry mimicking one half of the experimental patterned cell shape (Fig. 3A, S 2A-B).

**Figure 3:**
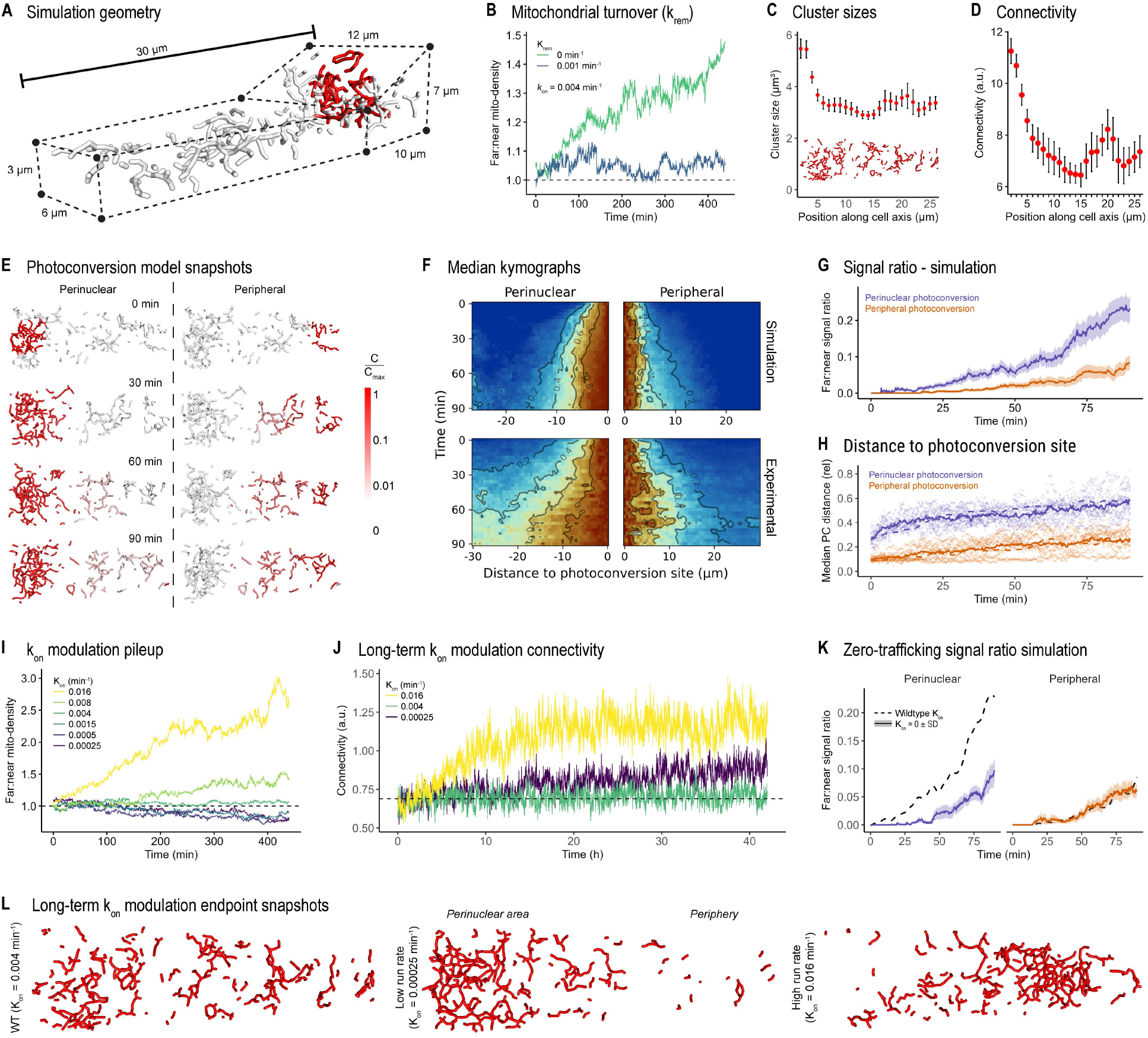
A simple physical model recapitulates many aspects of mitochondrial dynamics and mixing. (A) Simulation cell geometry; the dashed lines indicate the outer membrane of the simulated cell; the taller portion represents the perinuclear region with a photoconverted volume (red). (B) Wildtype-like anterograde transport *(rate constant k*_*on*_) causes mitochondrial peripheral pileup (green curve), which is balanced by the introduction of perinuclear growth and global turn-over *(rate constant k*_*rem*_), (blue curve). (C-D) Size and connectivity of mitochondrial clusters at wildtype-like parameters, recapitulating experimental observation of enhanced connectivity in perinuclear regions (N=20 simulations). (E) Snapshots of the mitochondrial network over time for simulated photoconversion experiments; mitochondrial segment color corresponds to the simulated photoconverted protein concentration. (F) Median ratio kymographs of photoconverted signal to mitochondrial mass along the long axis of the cell for each type of photoconversion; comparison between simulation (top) and experimental (bottom) results (experimental: perinuclear, N=29 and peripheral, N=16). (G) Integrated ratio of signal to mass (as in F), compared for positions less than or greater than 15 μm from photoconversion site. (H) Relative median distance to the photoconversion site for mitochondria containing photoconverted signal above threshold, as compared to all mitochondria. Dashed lines represent experimental averages (replotted from Fig. 1O). (I) Ratio of mitochondrial densities (as in B) for different run rates k_on_. (J) Long-term effects of k_on_ modulation on network connectivity (as in D). (K) Integrated ratio of signal to mass beyond 15 μm (as in G) without directed transport. Wildtype data as dashed lines. (L) Network morphology snapshots at the endpoint of long-term simulations, for three different values of k_on_.

To reflect our experimental observations of motor-driven mitochondrial motion, we incorporated biased transport directed toward the periphery for small mitochondria located in the perinuclear region. This motion is parametrized by kon, the rate constant for an individual mitochondrion to become associated with a motor protein attached to the cytoskeleton. Once in motion, mitochondria are transported outward over a distance drawn from an exponential distribution (average L/2, half the length of the cell arc) before returning to a diffusive state.

Setting the value of kon to achieve the experimentally observed run rate (~0.1 runs/min) (Fig. S 2C-D) resulted in the pileup of organelles at the cell periphery on the timescale of a few hours (Fig. 3B, green curve). Since peripheral mitochondrial accumulation was not observed experimentally, we hypothesized that biased anterograde transport exists to counteract an underlying asymmetry, such as perinuclear mitochondrial biogenesis. We thus introduced mitochondrial growth and turnover into the model, defining a rate krem for the spatially uniform removal of mitochondrial tips and fragments. Removal is a proxy for all processes that reduce mitochondrial density in the cell, including active degradation and dilution due to cell volume growth during the cell cycle, which would have been more complex to model explicitly. The same parameter also encompasses mitochondrial degradation (thought to occur at much slower timescales than the simulated events) (Kim et al., 2012). Production was, in turn, restricted to the perinuclear region, matching the removal events to maintain a constant overall mitochondrial density. The value of rate krem was chosen such that the entirety of the mitochondrial mass turns over in 42 hours, approximately matching the cell cycle time (Fig. S 2E-H). Incorporating directed runs and biased organelle biogenesis into the model resulted in a balanced distribution of mitochondrial mass between the peripheral and perinuclear regions (Fig. 3B, blue curve). Using the same parameter values, the model also replicated the experimentally observed spatial variation in mitochondrial size, with a perinuclear population presenting larger and more interconnected mitochondria (Fig. 3C-D). In our model, the enhanced perinuclear connectivity is a geometric consequence of the cell shape (Fig. S 2I), with larger clusters less likely to form in the narrow periphery due to excluded volume effects. These spatial differences in cluster size persist even in the absence of biogenesis and transport. Thus, our simple model including biased mitochondrial transport, growth, and balanced turnover, captures essential features of the experimental data.

To assess how the balance of biased transport and biased growth affects the spreading of material, we simulated a photoactivation experiment. Briefly, we activated a 36 µm^2^ patch of mitochondria in either the perinuclear or peripheral region of the simulated cell at time t=0. We then allowed the network to remodel according to the rules described above (Fig. 3E) and included a minimum threshold concentration (Methods) to account for experimental noise when generating simulated kymographs (Fig. 3F). While experimental kymographs show a broader mitochondrial density distribution at later timepoints, likely arising from heterogeneous cell and mitochondrial network sizes, the simulation closely replicates the differential spreading behavior between perinuclear and peripheral mitochondria measured in cells. We quantified the asymmetry in spreading by plotting the ratio of signals in a region far from versus near to the photoconversion site over time (Fig. 3G), finding that the ratio grows faster for photoconverted regions near the nucleus than those in the cell periphery. Furthermore, the relative median distance from the photoconversion zone for mitochondria containing above-threshold photoconverted signal increased more rapidly for perinuclear than for peripheral photoconversion (Fig. 3H), as was also observed in experimental data (Fig. 1K).

Since the model replicated key features of the experimental data, we used it to predict the effect of motor-driven transport on the mass distribution. First, to test the direct impact of asymmetric trafficking, we selectively removed the directed transport in the simulation while keeping all other empirical rates fixed. The spreading of photoconverted signal is thus only a consequence of mitochondrial fusion and growth. While peripheral signal spreading shows negligible change (Fig. 3K), the spreading of perinuclear signal is blunted down to peripheral levels, suggesting that the experimentally observed directed runs are critical to explain the bias in signal spread during photoconversion experiments, which cannot solely be explained by fusion and diffusion of signal throughout the interconnected network.

Increased anterograde run rates (high k_on_) resulted in mitochondrial accumulation at the cell periphery, while decreased rates (low k_on_) led to perinuclear pileup (Fig. 3I). Importantly, using experimentally derived values for k_on_ and k_rem_ (from running and growth rates) yielded an approximately uniform mass distribution (Fig. 3I), indicating that mitochondrial transport and growth rates in these cells are carefully balanced. Simulations over longer time periods (42 hours, comparable to 1 cell cycle) maintained this result, with increased or decreased run rates causing peripheral or perinuclear accumulation, respectively, and wildtype experimental run rates resulting in a stable morphology in time (Fig. 3J-L). Intriguingly, altering anterograde transport additionally resulted in an increase of overall model network connectivity over time (Fig. 3J), caused by the pileup of mitochondria promoting higher rates of fusion.

We note that the asymmetric spreading observed in the *in silico* simulations results from a combination of factors. First, biased anterograde transport enhances spreading from the perinuclear zone. Second, increased peripheral fragmentation (Fig. 3C-D) makes it more difficult for the signal to spread from the periphery. These effects encompass the interplay between mitochondrial dynamics (biogenesis, transport) and cellular geometry (increased perinuclear volumes), which governs both the spatial distribution of and the material exchange between mitochondrial clusters.

### Balanced motor-driven transport is required to maintain mitochondrial network dynamics and morphology

We sought to test the predictions of our model by experimentally measuring the impact of perturbing transport through the modulation of TRAK1 and TRAK2. The TRAK adaptors are anchored to mitochondria and mediate their bidirectional trafficking along microtubules (Glater et al., 2006; van Spronsen et al., 2013). While TRAK1-kinesin drives anterograde trafficking (away from the PNC), TRAK2-kinesin mediates anterograde, and TRAK2-dynein mediates retrograde transport in a conditional manner (Fenton et al., 2021).

Silencing TRAK1 expression with siRNA (TRAK1 KD) inhibited anterograde transport without affecting retrograde transport, consistent with reduced kinesin binding, while the overexpression (OE) of either adaptor enhanced retrograde transport (Fig. 4A-B, S 3A-C). None of our silencing and OE strategies boost anterograde transport, suggesting that baseline trafficking from the PNC is either at saturation or that adaptor recruitment is not a limiting factor. These results defy a simple model of adaptor abundance driving mitochondrial transport likelihood and instead hint at secondary impacts of transport, which in turn affect motility (*e*.*g*. larger mitochondria move less).

**Figure 4:**
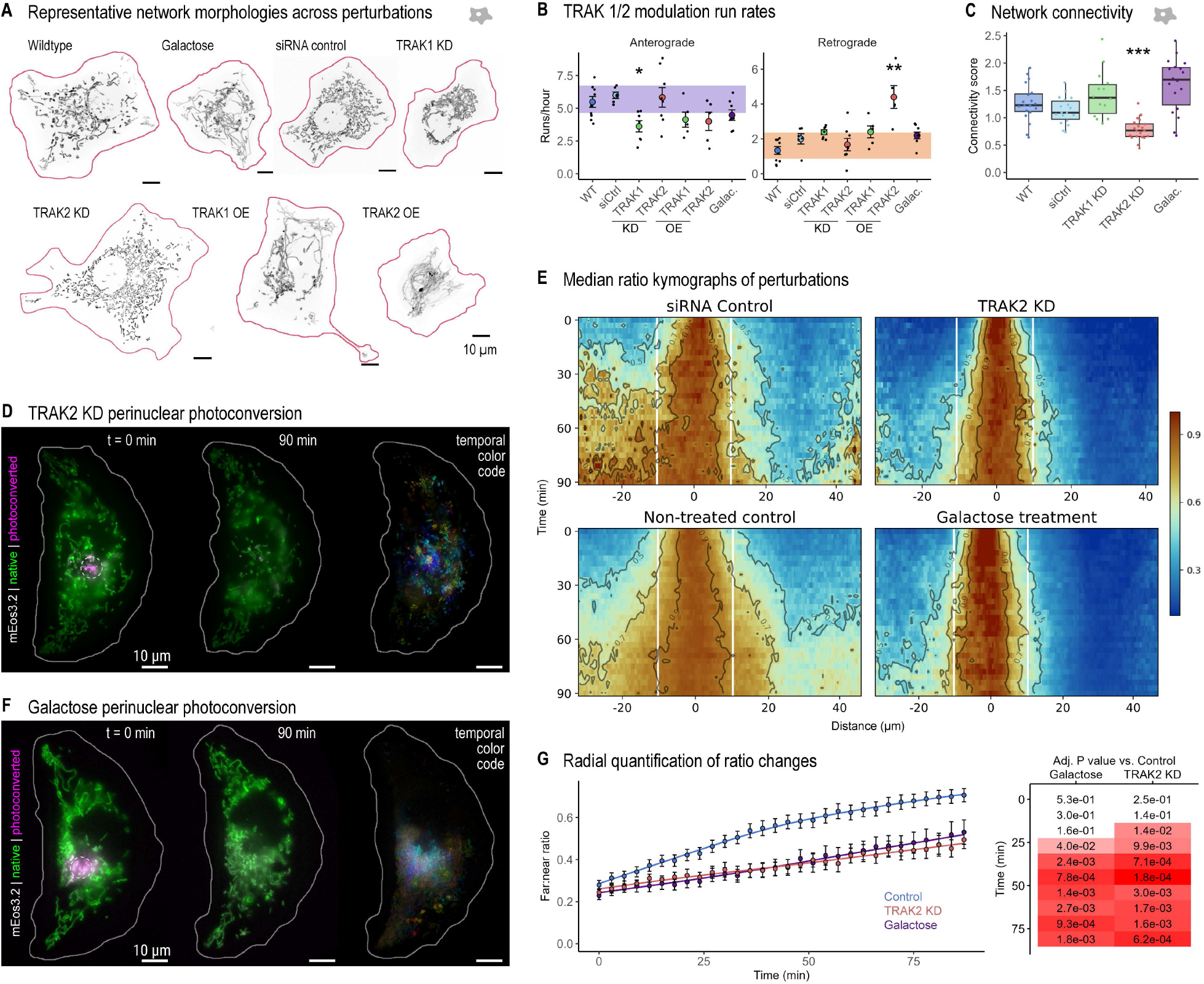
Modifying motor-driven transport or metabolism disrupts mass balance and connectivity. (A) Maximum intensity projections of representative mitochondrial networks resulting from different perturbations; mito-mScarlet signal (black), cell outline (red). (B) Quantification of transport rates for each TRAK perturbation condition represented as runs/min. N_TRAK1 KD_ = 7 cells, N_TRAK2 KD_ = 8 cells, N_TRAK1 OE_ = 5 cells, N_TRAK2 OE_ = 5 cells. Shaded area represents the SD of controls (non-treated and siRNA controls). (C) MitoGraph connectivity score for TRAK KD perturbations obtained from volumetric imaging; N_WT_ = 21, N_siCtrl_ = 18, N_TRAK1 KD_ = 15, N_TRAK2 KD_ = 20 cells. (D) Perinuclear photoconversion of a representative TRAK2 KD cell; merged native and photoconverted mEos3.2 (green and magenta respectively) at the start and end frames, as well as a temporal color-code of the photoconverted channel across the entire time series (right). Dotted circle, approximate photoconversion region. (E) Median of ratios kymograph for perinuclear photoconversions in siRNA control (N=14 cells), TRAK2 KD (N=19 cells), non-treated control (N=9 cells) or 16h-galactose treatment (N=18 cells). Horizontal axis spans the tip-to-tip cell extent, zero marks the site of photoconversion, vertical white lines indicate the 11 μm instantaneous diffusion boundary. (F) Representative perinuclear photoconversion of a cell following 16h-galactose treatment, as in (D). (G) Ratio between the median values beyond (far) and within (near) the instantaneous diffusion boundary as in (E), across perturbations. All scale bars 10 µm. WT = wildtype/non-treated control, siCtrl = siRNA control, KD = siRNA-mediated knock-down, OE = ectopic overexpression, galac. = Galactose treatment. Ameboid symbol indicates the data comes from non-empanada-micropatterned cells. P values from a Kruskal-Wallis/Dunn’s test with Bon-ferroni correction against controls (p< 0.05*, 0.01**, 0.001***).

Indeed, perturbations to TRAK had visible consequences on network morphology and interconnectivity (Fig. 4A). TRAK1 KD and the overexpression of either adaptor typically led to elongation and hyperfusion around the cellular center (Fig. S 3D). This is consistent with model predictions that reduced anterograde transport should lead to accumulation of mitochondria in the perinuclear region, increase encounter rates, and thereby increase connectivity (Fig. 3I-K, S 3C-D). TRAK2 KD had the opposite effect, displaying mitochondrial fragmentation without perinuclear clustering. Quantifying network structure confirmed a modest increase in network connectivity for TRAK1 KD and a decrease for TRAK2 KD compared to siRNA control cells (Fig. 4C).

Assessing the spread of material by photoconversion in the presence of TRAK1/TRAK2 perturbations was challenging, as most cells lacked peripheral mitochondria. As an exception, TRAK2 KD presented comparable network coverage to WT, albeit with a hyper-fragmented morphology. The fragmentation suggests that TRAK2 affects mitochondrial fusion, perhaps by mediating encounter rates. Despite the increased motility of smaller mitochondria and the unchanged run rates, TRAK2 KD presented a remarkably slower spread of perinuclear material (Fig. 4D-E), suggesting that reducing fusion rates can inhibit efficient material spreading despite wildtype-like active transport.

Stimulating oxidative phosphorylation by growing cells in glucose-deficient media supplemented with galactose leads to myriad metabolic and structural adaptations, which include the hyperfusion of the mitochondrial network (Gomes et al., 2011) (Fig. 4A). After 16h treatment the network extended across the cell, with increased connectivity score and a modest decrease in anterograde runs (Fig. 4B-C), in contrast to the increase in mitochondrial motility typically observed in neurons under similar conditions (Pekkurnaz et al., 2014). Unexpectedly, despite the hyperfusion, photoconverted perinuclear material spread more slowly than in control conditions (Fig. 4E-F). This indicates that network connectivity is insufficient to achieve material spreading across the entire cell, likely due to the lack of complex branched morphologies (Chuphal and Brown, 2024) and potential matrix compartmentalization by the inner membrane (Tábara et al., 2024; Malka et al., 2005).

Overall, our data qualitatively replicate the predictions of the *in silico* model and indicate that modulating microtubule-based transport impacts the relative rates of anterograde and retrograde trafficking, which drives changes in network morphology that in turn feed back on motility. Material spreading beyond the PNC was similarly reduced in both TRAK2 KD and galactose treatment, despite comparable network coverage to WT and opposite morphological changes (fragmentation and hyperfusion, respectively). Hyperfused networks allow for effective spreading of material over short distances but are incapable of supplying mitochondria throughout the whole cell. In contrast, hyperfragmented networks can support motility of individual organelles, but lacking fusion-based content sharing may not allow for wild-type levels of material mixing.

In summary, mitochondrial transport has been systematically studied in the axons and dendrites of neurons, but the study of mitochondrial transport in unpolarized cell types has remained limited – relying on indirect measures such as snapshots of the mitochondrial network and its extent in the cell (López-Doménech et al., 2018), or on measurements in the cell periphery (Mills et al., 2016). Our data reveal that cells lacking polarized function can present spatially organized, functionally distinct mitochondrial populations and biased trafficking.

Mitochondrial biogenesis requires the orchestrated expression of nuclear and mitochondrial genomes to ensure the correct import, folding, and assembly of the ~1300 proteins comprising the mitochondrial proteome (Rath et al., 2021). Biogenesis has been previously proposed to occur in the perinuclear region (Davis and Clayton, 1996), which could offer a fitness advantage by increasing the availability of nuclear transcripts and deoxynucleosides needed for mtDNA replication. The location of mtDNA replication has been studied in cultured cells, but results have remained inconclusive (Davis and Clayton, 1996; Magnusson et al., 2003), possibly due to DNA denaturation requirements and other limitations of traditional BrdU measurements. Our measurements highlight a preferential biogenic role for the perinuclear mitochondrial subpopulation. Rapid content mixing within this proliferative PNC can ensure a high degree of homogeneity among mitochondria, which are then supplied to other parts of the cell.

We find that mitochondrial material mostly spreads in the anterograde direction from the perinuclear population. Our spatially resolved dynamic model supports the notion that this asymmetric transport ensures material mixing while counter-balancing biased perinuclear biogenesis, avoiding the buildup of mitochondria in the center or periphery of the cell. Biased transport was previously reported in mouse fibroblasts, where peripheral motile mitochondria had a three-fold greater rate of anterograde versus retrograde runs (Mills et al., 2016). Thus, our study connects two phenomena that have been reported in a broad range of cell types: a subpopulation where biogenesis takes place, and redistribution via directed mitochondrial transport, establishing a spatially resolved model for the first half of a mitochondrial life cycle.

The role of motor-based motility in establishing the subcellular distribution of mitochondria has been well-documented, but its contribution to network morphology is rarely described. TRAK1 has been suggested to affect fusion and connectivity in HeLa cells (Lee et al., 2018), where its absence leads to perinuclear accumulation (Barel et al., 2017, 20). We observe that modulating TRAK1 and TRAK2 in COS-7 cells can have major implications for network morphology. While some of these effects could be expected from transport-related changes in encounter rates, others may arise from alterations in the duration or directionality of the forces that motors apply to organelles, leading to differences in fission and fusion rates or pulling of thinner tubules. Intriguingly, TRAK1 and TRAK2 can present tissue-specific expression (Uhlen et al., 2010), or even polarized distribution within neurons (van Spronsen et al., 2013), while TRAK1 mutations are associated with reduced mitochondrial membrane potential and lethal encephalopathy (Barel et al., 2017). Whether and how defects in coupling mitochondria to molecular motors impact mitochondrial quality control or overall metabolic function remains to be investigated.

Taken together, our study highlights the presence of asymmetries in mitochondrial populations and the interdependence between microtubule-based trafficking and network morphology in content mixing. While fission and fusion are well-known modulators of mitochondrial morphology and dynamics, trafficking should also be considered a key player. Finally, differences between mitochondrial subpopulations should be considered, for example, in microscopy studies which may focus on peripheral populations that are easier to image.

## Data availability

The numerical data underlying the figures are openly available in Zenodo (accession number: 19631627). Large imaging data are available from the corresponding author upon reasonable request.

## Acknowledgements

We thank Hélène Perreten Lambert for her support with cell culture, cloning, and western blotting and we thank Kyle Douglass for his coding support and rich feedback on data and image analysis. We would also like current and past members of the laboratory of experimental biophysics for their help and feedback in the development of this project. Research in S.M.’s laboratory is funded by the Swiss National Science Foundation (SNSF) Grant 10268 and the ERC Consolidator Grant Piko. EFK and KH were supported by the US National Science Foundation, grant #2310229, and the Chan Zuckerberg Initiative.

The authors declare no competing financial interests

## Author contributions

Conceptualization JW, JCL, SM

Methodology EK, SM JW, JCL, KH,

Investigation SBN, EBD, TK JW, JCL, KH,

Visualization JW, JCL, KH

Writing – original draft JW, SM, JCL

Writing – review and editing All authors

Supervision SM

Project administration SM

Funding acquisition SM

## Materials and Methods

### Plasmids and reagents

The following plasmids were used. pMTS_mScarlet_N1 (mito-mScarlet) was a gift from Dorus Gadella (Addgene plasmid #85057), mito–GFP (Cox-8 presequence) was a gift from H. Shroff (NIH, Bethesda, USA), mito-StayGold (pCSII-EF/mt-(n1)StayGold) was a gift from Atsushi Miyawaki (Addgene plasmid #185823), pRSET-mEos3.2 was a gift from T. Xu (Chinese Academy of Sciences, Beijing, China), GFP-Trak1 was a gift from Josef Kittler (Addgene plasmid #127621), and GFP-Trak2 was a gift from Josef Kittler (Addgene plasmid #127622). Mito-mEos3.2 was amplified from mito-GFP and pRSET-mEos3.2. TRAK1-mito-mEos3.2 and TRAK2-mito-mEos3.2 were amplified from TRAK1/2-GFP and mito-mEos3.2 using the P2A self-cleaving peptide system.

Oligonucleotides for siRNA were designed with and made by Microsynth. *TRAK1*(sense strand 5’-GAA GCU GAA AGA CCU UGA A dTdT-3’) and *TRAK2* (sense strand 5’-UCA GAG UCU UCC AGU CAU A dTdT-3’). Cells were incubated with 12.5pmole of siRNA for 48h before seeding on patterned coverslips for imaging the subsequent day.

### Cell culture, transfection and seeding

COS7 cells were grown in Dulbecco’s modified Eagle’s medium (DMEM) supplemented with 10% fetal bovine serum and maintained in culture for a maximum of 20 passages. For galactose medium, glucose-free DMEM was supplemented with 10% dialyzed FBS (Thermofisher, A3382001). Cell medium was changed for galactose medium either immediately upon transfection (approx. 48h before imaging) or upon cell seeding on the patterned coverslips (approx. 16h-22h before imaging).

Transfection for wt imaging of mito-mEos3.2 was performed in a T25 flask split at 1/5^th^ dilution from the main flask 24h before transfection. All other plasmid transfections were performed 24h after plating cells in a 6-well plate at 40’000 cells well^-1^. Plasmid transfections were performed with Lipofectamine 2000 (Invitrogen) in Opti-MEM medium (Invitrogen) following the manufacturer’s instructions for 6-well plates. For transfections in T25 flasks, the protocol for one well was tripled. For photoconversion experiments with siRNA, the siRNA was co-transfected with mito-mEos3.2 with lipofectamine 2000. For mitochondrial volume imaging, siRNA was transfected using lipofectamine RNAiMAX (Invitrogen) 24h after plating and mito-mScarlet was transfected another 24h later with lipofectamine 2000. In both cases the cells were not re-transfected upon seeding the cells on patterned coverslips the night before imaging.

Cells were seeded on micropatterned coverslips the evening before imaging (approx. 16h) at 20’000 cells coverslip. To ensure a homogeneous distribution, the cells are added dropwise and left to settle for 5 minutes before moving to the incubator. After 30-60 minutes the cells are washed once with fresh medium and dishes are filled with 2ml of phenol-red-free medium. For imaging experiments on unpatterned coverslips, cells are plated on 25mm coverslips prior to transfection and remain untouched until imaging.

Nocodazole treatment was performed using a 5 µM concentration, prior to which cells labeled with 500 nM MitoTracker and 1x SPY555-Tubulin (Spirochrome, following manufacturer’s instructions), and imaged for 5 frames at 1 min time intervals. After 1 hour of Nocodazole treatment, the same cells were imaged with identical conditions. Nocodazole effect was confirmed by the tubulin phenotype. For quantification of mitochondrial displacement, a published mitochondrial motility measurement analysis pipeline was adapted (Han et al., 2023). In brief, mitochondrial motility was quantified by calculating the displacement (travel area) for subsequent frames relative to the previous frame.

#### Micropatterning

The process for making patterned coverslips tightly follows the protocol published by *Azioune et al*. (Azioune et al., 2010).

Patterns were designed to fit a standard 4” quartz glass chrome blank using the Auto-CAD software from Autodesk. The designs were then converted to the.gds format for laser writing to a 4” chrome blank in-house at the Center for MicroNanoTechnology (CMi) at EPFL using a Heidelberg Instruments VPG200.

The glass bottom of 35mm culture dishes (MatTek, P35G-1.5-14-C) was removed with a scalpel and the patterned coverslip was glued in place (Loctite, SI 5145). Immediately before seeding cells, the patterns are coated with fibronectin using 300 µl of coating solution containing 25 µg/ml of fibronectin (Sigma-Aldrich, F1141-1MG) in 100mM NaHCO_3_ pH8.5.

### Immunoblotting & immunofluorescence

#### Immunoblotting

For immunoblots 48 hours after siRNA transfection, COS-7 cells were lysed in RIPA buffer (Sigma) supplemented with fresh proteases inhibitors (Sigma Aldrich 11836170001) on ice for 30 minutes. A centrifugation at 16,000 g for 10 min at 4°C was performed to remove the insoluble material. Protein concentrations were determined using a Pierce BCA protein assay kit (Life Technologies, 23227) and equal amounts of protein were analyzed by self-casted 12.5% SDS-PAGE (30-50 μg of proteins per lane). For immunoblotting, proteins were transferred to nitrocellulose membranes (BioRad) electrophoretically and incubated with the specified primary antibodies (see above), diluted in 5% non-fat dry milk in Tris buffered saline with Tween 20 (TBST). The blots were further incubated with anti-rabbit HRP conjugated secondary antibodies (GE Healthcare) and visualized using ECL (GE Healthcare). The following primary antibodies were used for immunoblotting: anti-TRAK1 (Invitrogen, PA5-44180, 0.5 µg/ml) and anti-TRAK2 (Proteintech, 13770-1-AP-150UL, 1:500 dilution)

#### Immunofluorescence

Fixation was performed in pre-warmed 4% PFA in PBS for 15min, followed by 3 washes in PBS. Permeabilization and blocking were performed in a single step by incubating the fixed cells in 10% pre-immune goat serum (Thermo Fisher), 0.3% Triton X-100 (Sigma) in PBS for 30min. The same buffer was used to dilute the specific primary antibodies (anti-TRAK1 (Invitrogen PA5-44180, 1:250), anti-TRAK2 (Proteintech 13770-1-AP, 1:250)). After 2h of incubation, cells were washed in PBS and incubated for 1h in the appropriate secondary antibody (Anti-rabbit Alex Fluor 488 (Invitrogen A-11034, 1:500). Nuclei were then stained by incubation with Hoechst for 15 min after secondary antibody incubation. To quantify TRAK1 and TRAK2 amounts upon siRNA silencing, blinded images were analyzed using the CellProfiler software: Shortly, nuclei were segmented and used to identify and segment each individual cell’s cytoplasm by propagation and masking from the inverted phase-contrast image (using the IdentifyTertiaryObjects module). Then, the mean intensity values were obtained for each cell using the MeasureObjectIntensity module on the immunofluorescent channel.

### Microscopy

#### Widefield imaging

Imaging of nocodazole treated cells and imaging of photoconvertible mito-mEos3.2 was done on a Zeiss Axio Observer 7 inverted microscope equipped with an sCMOS camera (Photometrics Prime), CoolLED pE-800 LED illumination system and 405nm FRAP system (VisiFRAP, Visitron). The imaging objective was a 63x 1.40 NA oil immersion objective (Zeiss Plan-Apochromat 63/1.3 Oil Ph3 M27).

#### Photoconversion

The photoconversion samples were first thoroughly searched for properly patterned cells with high green fluorescence and saved for multi-point acquisition. Between 10-18 cells are imaged in each acquisition. Before launching the acquisition, a circular ROI with a diameter of 7 µm is placed at the MTOC or the tip of the “arm” of the cell on the side closer to the PNC (so the nucleus does not obstruct transport). The ROI was then photoconverted and the process was repeated for each cell of the set. This process created a delay of approx. 3 minutes between photoconverting the first cell and launching the acquisition (1 min/5 cells). Generally, peripheral mitochondria were only photoconverted if there was significant peripheral population. Cells with no or very few peripheral mitochondria were not considered. Images were captured in the following sequence: phasecontrast, 470 nm excitation (2%), 550nm (20%) and emission was captured with dedicated filter cubes for GFP and Texas-red. For 90-minute acquisitions the intervals were set to 3 minutes and for high-speed acquisitions images were captured every 2 seconds omitting the phase-contrast channel.

All imaging was done at 37°C and 5% CO_2_ in phenol-red-free DMEM. The microscope was controlled by the imaging software VisiView 5.0 from Visitron.

#### Confocal imaging

Imaging of RNA FISH, EdU, and volumetric imaging of the mitochondrial network were performed on a Nikon Eclipse Ti2-E inverted microscope equipped with an sCMOS camera (Photometrics Prime 95B) and a 100x 1.49 NA oil immersion objective (SR HP Apo TIRF 100x/1.49, Nikon).

### Image Analysis

#### Mitochondrial volumetric analysis

Z-stacks of cells labelled with mito-mScarlet, excited with a 561nm laser at 80% power and exposure time of 20ms, were acquired with 100nm z-steps. For TRAK1 and TRAK2 overexpressions, TRAK1/2-GFP signal was checked for each cell to ensure appropriate and comparable overexpression. The imaging was done at 37°C and 5% CO_2_ in phenolred-free DMEM and the microscope was controlled by the NIS-Elements software from Nikon. Before further processing, the RAW images were cropped and deconvolved with Huygens Core offered by the EPFL imaging core facility (BIOP).

The deconvolved z-stacks are processed by mitograph (Viana et al., 2015) running via the Windows subsystem for Linux (WSL2). Tiff stacks are processed with the following command:./MitoGraph -xy 0.11 -z 0.1 -labels_off -path <file-path>. Once processed, the corresponding R-scripts from the mitograph authors aggregate the data and measurements into summary files which were used for plotting in python.

Comparison of mitochondrial volume to the distance to the PNC was done by opening the output mitosurface.vtk files and determining the center of mass for each connected component (mitochondrion) and the total center of mass of all mitochondria (PNC). The wildtype control 3D volumetric data of mitochondrial networks on patterned surfaces was additionally utilized to optimize the mitochondrial network architecture model in (Holt et al., 2024).

#### Fission quantification

Timelapses of the mitochondrial network were obtained from unpatterned COS-7 cells transiently expressing mito-StayGold. Fission events were manually detected and classified as “midzone” when the smaller daughter is at least 25% of the full-length larger daughter mitochondrion, and otherwise “peripheral”. Next, the distances between each fission and the PNC (manually approximated as the center of the cell and following the radiation of the mitochondrial network) were measured blindly to fission type.

#### Kymograph generation and analysis

Following are the steps to generate average kymographs from the mito-mEos3.2 timeseries.

#### Line profile generation and processing

Timelapse movies are opened in Fiji and a segmented line ROI with a width of 100px is defined along the long axis of the cell in the phase contrast channel. A custom macro then measures and saves the average intensity along the ROI for the photoconverted and unphotoconverted channels.

The individual line profile files are further processed in python. First the line profiles are binned by a factor of 5, then the photoconversion point is determined and the distance values are centered around this point. Finally, the ratio of the photoconverted to the unphotoconverted signal is calculated. Photoconverted, unphotoconverted and ratio values are saved as individual columns in a tidy dataframe as a.csv file.

#### Kymograph generation and averaging

The tidy dataframes are now combined by either perinuclear or peripheral photoconversion, centered on the photoconversion site with respect to each other and cropped to the smallest cells size. A mean kymograph is generated for the photoconverted and unphotoconverted channels, while a median kymograph is generated for the ratio channel to account for severe outliers from noisy data at the edges of the kymographs. The median kymographs displayed and used for analyses were calculated using unscaled signals, as ratio = native / (native + photoconverted). The full-width half-maxima (FWHM) of the photoconversion zone were calculated from the photoconverted signal peak at time = 0. The out:in ratios used for signal dissemination were calculated using a threshold of 11 μm, based on the median size of the largest interconnected mitochondria and median FWHM of instantaneous photoconversion (10.3 ± 3.1 μm and 11.6 ± 4.5 μm respectively).

#### Single mitochondria analysis

To further analyze the dynamics on the single mitochondrial level, the widefield mito-mEos3.2 images (photoconverted and unphotoconverted channels) were segmented using Ilastik (Berg et al., 2019). Following pre-processing (Fiji default subtract background command with a rolling-ball radius of 10px and median filter of size 2px), images were batch thresholded with a manually trained Ilastik algorithm. Using the Ilastik threshold as a mask, the data was further analyzed using the CellProfiler software (Stirling et al., 2021), identifying individual mitochondria using the IdentifyPrimaryObjects module and measuring their size, shape and distance to the photoconversion spot (computed as a 3.5 µm radius circle from the photoconverted channel’s maximum intensity pixel).

The data is then plotted as the median distance of the photoconverted mitochondria to the photoconversion spot of each cell normalized to that of non-photoconverted mitochondria of that cell.

#### Mitochondrial tracking

Mitochondrial tracking was performed manually using the manual tracking plugin for Fiji from Fabrice P. Cordelières. We focused on short bursts of motion over full tracking of a mitochondrion and classified each track by its direction (anterograde/retrograde) for further quantification of trafficking dynamics. These tracks were generated from movies acquired at high frame rates with an interval of 2 seconds.

To quantify the number of anterograde and retrograde moving mitochondria, all directedly moving mitochondria were quantified in multiple, randomly chosen movies from different imaging days from the datasets for kymograph generation.

### Transported volume estimation

Mitochondrial volume transported per time was calculated using the average run rate and the mean length of a mitochondrion undergoing directed motion, assuming mitochondria take on a cylindrical shape with a radius of 250 nm.

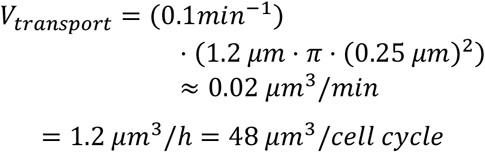

In turn, median mitochondrial volume in the periphery was calculated as: median network volume * fraction of mitochondrial network in periphery; this used the mean total length of a WT mitochondrial network, and assumed mitochondria take on a cylindrical shape with a radius of 250 nm. To find the peripheral fraction, we took the ratio between the integrated intensity of non-photoconverted mito-mEos3.2 for “radial class C,” corresponding to >22 μm from the PNC and the total intensity.

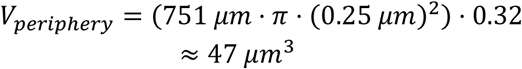

### EdU mtDNA replications assay

Replicating nucleoids were labeled using the Click-iT™ EdU Alexa Fluor™ 488 Imaging Kit (Fisher Scientific), following a modified version of the manufacturer’s instructions. Specifically, non-confluent cells were incubated in 50 μM EdU-containing media and fixed at different timepoints (15, 30, 45 and 60 min). Additional modifications from the protocol included two sequential Click reactions, and subsequent immunostained with a dsDNA antibody (abcam, ab27156) to label all nucleoids.

EdU labelled cells were imaged on the spinning-disk confocal microscope described above, with the 488nm and 561nm laser at 100% sequentially with an exposure time of 50ms. The z-step size was 200nm.

Images were analyzed using the CellProfiler software with manual curation. Shortly, following manual cropping of individual cells, nuclei were segmented from the dsDNA signal using the IdentifyPrimaryObjects function with a large size threshold. Then, the nucleus was masked from the dsDNA signal, and nucleoid foci were segmented using gaussian filtering and Robust Background algorithm with 2 standard deviations. Distances between nucleoids to the nucleus were automatically quantified, as well as each nucleoid’s EdU intensity. To classify the nucleoids, the EdU integrated intensity was normalized to the mean intensity of all nucleoids in that cell and used a threshold of 1. Other methods of classification, such as segmentation of EdU foci and classification by object overlap, were tested and generated comparable results.

### RNA FISH assay

Custom Stellaris® FISH Probes were designed against NDUFA10 by utilizing the Stellaris® RNA FISH Probe Designer (Biosearch Technologies, Inc., Petaluma, CA) available online at www.biosearchtech.com/stellarisdesigner (Version 4.2). The cells were hybridized with the NDFUA10 Stellaris RNA FISH probe set labelled with Quasar 670 (Biosearch Technologies), following the manufacturer’s protocol for simultaneous immunofluorescence and RNA FISH at www.biosearchtech.com/stellaris-protocols. For the probe and control, 30’000 cells were plated on 25mm coverslips 48h before performing RNA FISH. Cells were coincubated with the primary antibody for anti-TOMM20 (Abcam, ab186734, 1:1000 dilution) for 6 hours during the hybridization step of the protocol. Cells were incubated with a Goat anti-Rabbit IgG Alexa Fluor 488 secondary antibody (Invitrogen, A-11034, 2.5 µg/ml) during the wash stages, as well as DAPI during the last wash step. Instead of aspirating liquid from the wells we transferred the coverslips between wells of fresh washing buffer.

FISH labelled cells were imaged on the spinning-disk confocal microscope described above. Quasar 670 was imaged with a 638nm laser at 100% power and exposure time of 2s. Alexa 488 was excited with a 488nm laser at 80% power and 20ms exposure time. The z-step size was 200nm.

To analyze the images, the bigfish library (Imbert et al., 2021) for python was used. In brief the pipeline looked like this: Manually draw the cell outline in napari (napari, 2023), use the Li threshold to segment mitochondria, use the bigfish.segmentation unet_3_classes_nuc algorithm to segment the nucleus, detect spots with a spot radius of 150 x 150nm and apply subpixel fitting, classify the location of the spots with the segmentations. Random points were generated over the area of the bounding box of the cell outline for comparison with the real distribution of FISH foci.

### *In silico* modelling of mitochondrial dynamics

#### Model overview

Simulations of mitochondrial networks are carried out via a hybrid Brownian Dynamics and Monte Carlo approach, with individual discrete mitochondrial units driven by a random Brownian force, and potential gradients associated with bending, stretching, steric exclusion, and confinement within the cell. Mitochondrial units in proximity to each other are allowed to undergo tip-tip and tip-side fusion with an angle-dependent fusion rate and junctions can break apart with a constant fission rate. Details of the simulation methods are described in (Holt et al., 2024). In addition to the previously developed basic model, directed anterograde movement and dispersion of mitochondrial material within the network is incorporated in the current simulations as described below.

#### Half-cell geometry

Simulations are carried out in a domain with geometry illustrated in Figure 3a. This geometry is intended to represent half of a patterned COS-7 cell, with the 10 µm-long thicker region (0 < *x* < 10) corresponding to the zone near the nucleus on the side containing the PNC. The thinner region of the domain extends to the cell periphery, to 30-40 µm. For both cell geometries, the parameters are set to match experimental measurements as described below. The slightly different cell lengths are not expected to affect the conclusions drawn from the model study.

#### Selection of fusion parameters

To quantify the connectivity of experimentally observed and simulated networks, we use the global association constants *C*_*1*_, *C*_*2*_ for tip-tip and tip-side fusion, respectively defined as:

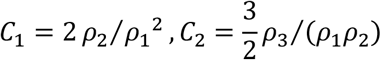

where *ρ*_*i*_ is the density of nodes of degree *i*. Degree-2 nodes are defined at 0.5 µm intervals along experimentally extracted network structures. For a total mitochondrial unit density ρ, values of 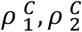 above 1 correspond to largely fused mitochondrial tips and junctions, respectively. Details of association constant calculations are described in (Holt et al., 2024). The fission rate is fixed to *k*_*f*_ = 0.5 min^-1^ at degree-2 junctions and 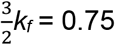 min^-1^ at degree-3 junctions. Fusion rates *k*_*u1*_, *k*_*u2*_ are obtained by a simulation parameter sweep, selecting the values that best match the observed association constants 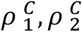 for mitochondrial networks in patterned COS-7 cells. Interpolating the simulation results gives predicted experimentally-relevant fusion parameters *k*_*u1*_ ≈ 700 min^-1^, *k*_*u2*_ ≈ 230 min^-1^ for wildtype cells in both the 30 µm and 40 µm cell geometries. We note that the smaller cell size also uses fewer mitochondrial units (550 vs 600) in order to keep a consistent density.

#### Directed runs

We model biased active transport of mitochondria along microtubules by enabling small mitochondria originating near the nucleus to engage in processive anterograde motion. Mitochondrial clusters containing no three-way junctions, of total length below 1.5 µm, and located within the thicker perinuclear region of the cell are capable of engaging in runs. Each such mitochondrion has a rate constant *k*_*on*_ for initiating a processive run. Once running, a mitochondrial unit is pulled by a harmonic spring (representing a motor) whose end-point moves with a constant velocity *v* = 15 - 20 µm/min along the long axis of the cell. The processive motion stops for one of three reasons. First, the unit dissociates from the motor at rate *k*_*off*_ = 2*v*/*L*, where *L* is the cell length. Second, a run ends whenever the force on the motor exceeds a stall force *F*_*stall*_, which may happen if a segment becomes tangled with other mitochondria in its way. Third, mitochondria stop moving processively upon reaching the boundary of the cell at the peripheral tip. We choose a value for *k*_*on*_ by matching the observed number of directed runs in simulations to that observed in experiment. This gives *k*_*on*_ = 0.0024, 0.0040 min^-1^ for 30 µm and 40 µm cells, respectively.

#### Turnover and growth

We model biogenesis of new mitochondria and accompanying cell growth via the birth of new mitochondrial units in the perinuclear region. To accomodate this growth without having to grow our cell boundary throughout the simulation, we pick terminal mitochondrial units (edges with at least one degree-1 node) to remove from the domain at random with rate *k*_*rem*_. Whenever a unit is removed, this triggers the birth of a new unit in the perinuclear region, where an edge is picked at random before growing rapidly until it reaches a length of 2*l*_*0*_, at which point the edge becomes two edges of unit length *l*_*0*_. We choose a value for *k*_*rem*_ by extrapolating the time to synthesize the full mitochondrial mass in simulations from the measured rate of new edge growth events. This time is then matched to the whole-cell doubling time of COS-7 cells (35-48 hours) (DSMZ-German Collection of Microorganisms and Cell Cultures GmbH, Leibnitz Institute), with *k*_*rem*_ chosen to give the closest agreement. This procedure gives *k*_*rem*_ = 0.0012, 0.0010 min^-1^ for 30 µm and 40 µm cells, respectively (Fig. S2 E-F).

#### Simulations of photoactivated material spread

To model the dispersal of photoconverted material, similar to the experimental data in Fig. 1, we use a Finite Volume Method approach to propagate diffusing photoactivated proteins across connected mitochondrial units. At each junction, the concentrations of the connected edges are updated according to:

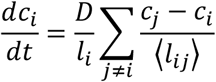

Where the *c* terms are the concentrations of protein on each edge, the ⟨*l*⟩ terms are the average edge-lengths of the relevant mitochondrial units, and *D* is the effective protein diffusivity. We use a protein diffusivity of *D* = 50 µm^2^/min.

A region of size 6 µm * 6 µm, projected downward along the z-axis is `photoconverted’ at the start of the simulation, with each mitochondrial unit overlapping with that region acquiring a normalized concentration of *c*_*i*_ = 1. The signal then propagates through fused mitochondrial units over time.

#### Analysis of fluorescent signal spread in simulated mitochondria

Because experimental images include noise and background signal absent in simulated data, we assume there is a nonlinear relationship between apparent fluorescence and photoconverted protein concentrations in the simulated mitochondria.

First, we assume that some minimal protein concentration is necessary in order for any photoconverted signal to be visible in a mitochondrial unit. This threshold concentration (*c\**) is substantiated by considering the median distance of mitochondria visible (labeled with photoconverted signal) versus not, and comparing to experimental data. We follow the same procedure outlined for experimental data (Fig. 1O) to calculate the median relative distance to the photoconversion site. To translate from our half-cell to the full cell geometry, we reflect the positions of all mitochondria across the *x* = 0 plane, assuming that no signal penetrates into this reflected region. We ignore any effect of the nucleus on mitochondrial positioning. A threshold concentration set to 2.9% of the maximum photoconverted signal per mitochondrial unit is found to give the best fit for the perinuclear activated region. The same threshold concentration also reproduces the slower increase in relative median distance from the peripheral photoconverted region.

Next, we consider in the experimental data individual mitochondria far from the site of photoconversion that have some visible signal in the photoconverted channel during the first frame of the imaging. On average, after subtracting global background, the signal in these mitochondria is *c*_*vis*_ = 0.15, as a fraction of the maximum photoconverted signal. This number is taken to represent an additional fluorescent background signal specifically localized to mitochondria. In analyzing the simulations, we thus take the effective fluorescent signal to be defined by the step-wise function incorporating both a threshold for visibility and a per-mitochondrion background: *f*_*i*_ = 0 if *c*_*i*_ < *c*, f*_*i*_ = 0.15 if *c\** < *c*_*i*_ < *c*_*vis*_, and *f*_*i*_ = *c*_*i*_ if *c*_*i*_ > *c*_*vis*_.

#### Generation of kymographs

To generate the kymographs, we first create binned line profiles along the long axis of the cell for photoconverted and background signal. At each timestep, the simulation cell is partitioned into bins of width 0.5 µm. Each mitochondrion is assigned an effective fluorescent intensity *f*_*i*_ as described above. The sum of these intensity values in each spatial bin represents the ‘red’ (photoconverted) signal measured in experiment, while the total number of mitochondria in each bin represents the ‘green’ signal. We then calculate the ratio of red to green signal at each timepoint and each bin position before averaging across 10 simulation trials (corresponding to 10 independent iterations with identical parameters). Finally, the resulting kymographs are cropped in time and space to 1 – 91 min post-photoactivation and 0 – 27 µm (0 – 37 µm for 40 µm cells) from the center of the photoactivation region. Lastly, we subtract the global (all time and space) minimum ratio and normalize by the maximum ratio at each timepoint.

## Supplementary figures

**Figure S1:**
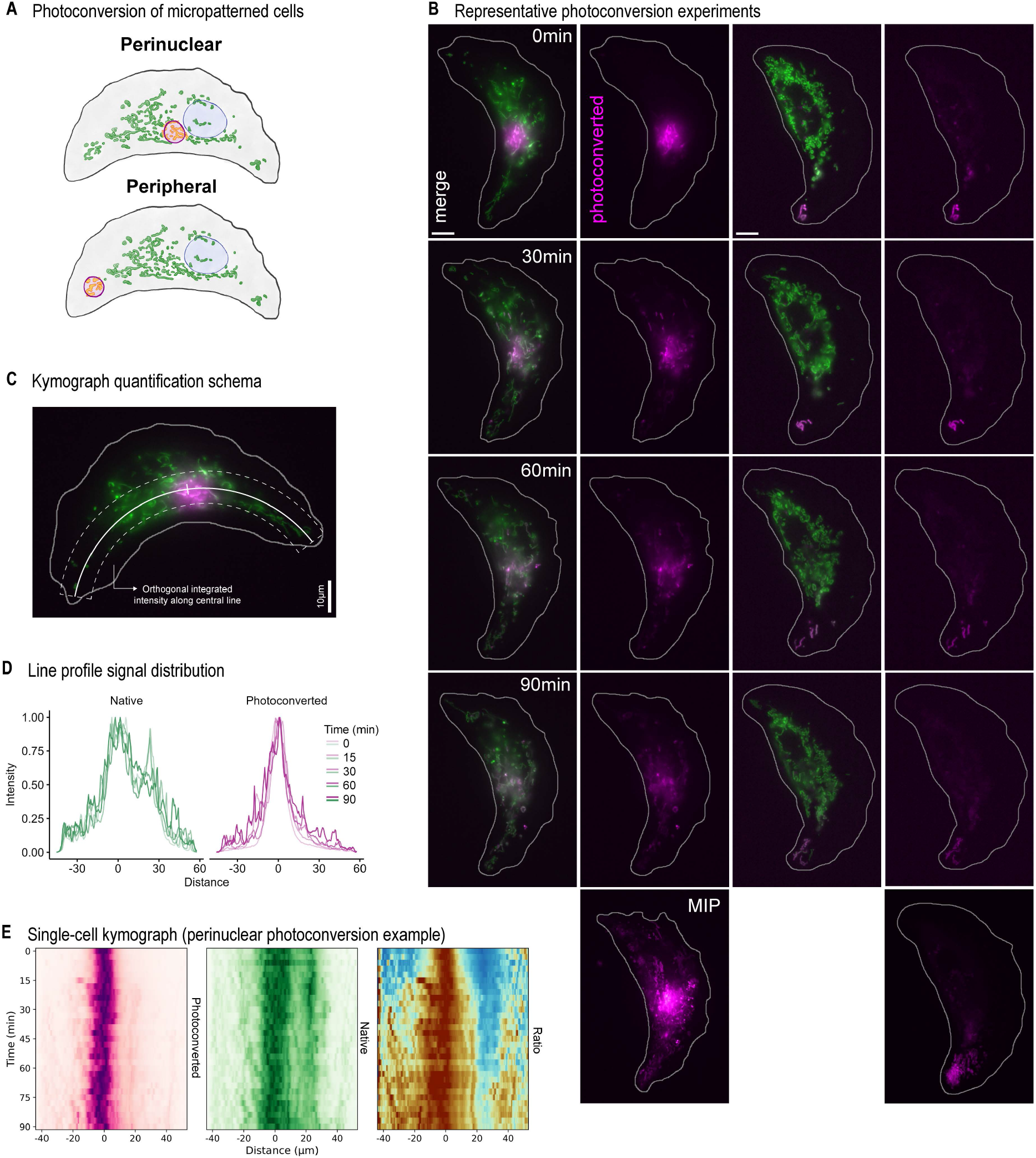
(A) Graphical depiction of the different photoconversion approaches. (B) Snapshots of two representative photoconversions as merged and photoconverted channel only (magenta). Maximum intensity projection (MIP) of the photoconverted channel (bottom). (C-D) Quantification framework for kymograph generation: representative thick line positioning (C) and resulting normalized line profiles along the cellular long axis for each timepoint (D). (E) Single-cell kymographs resulting from (E), for each channel and photoconverted:native ratio (right).

**Figure S2:**
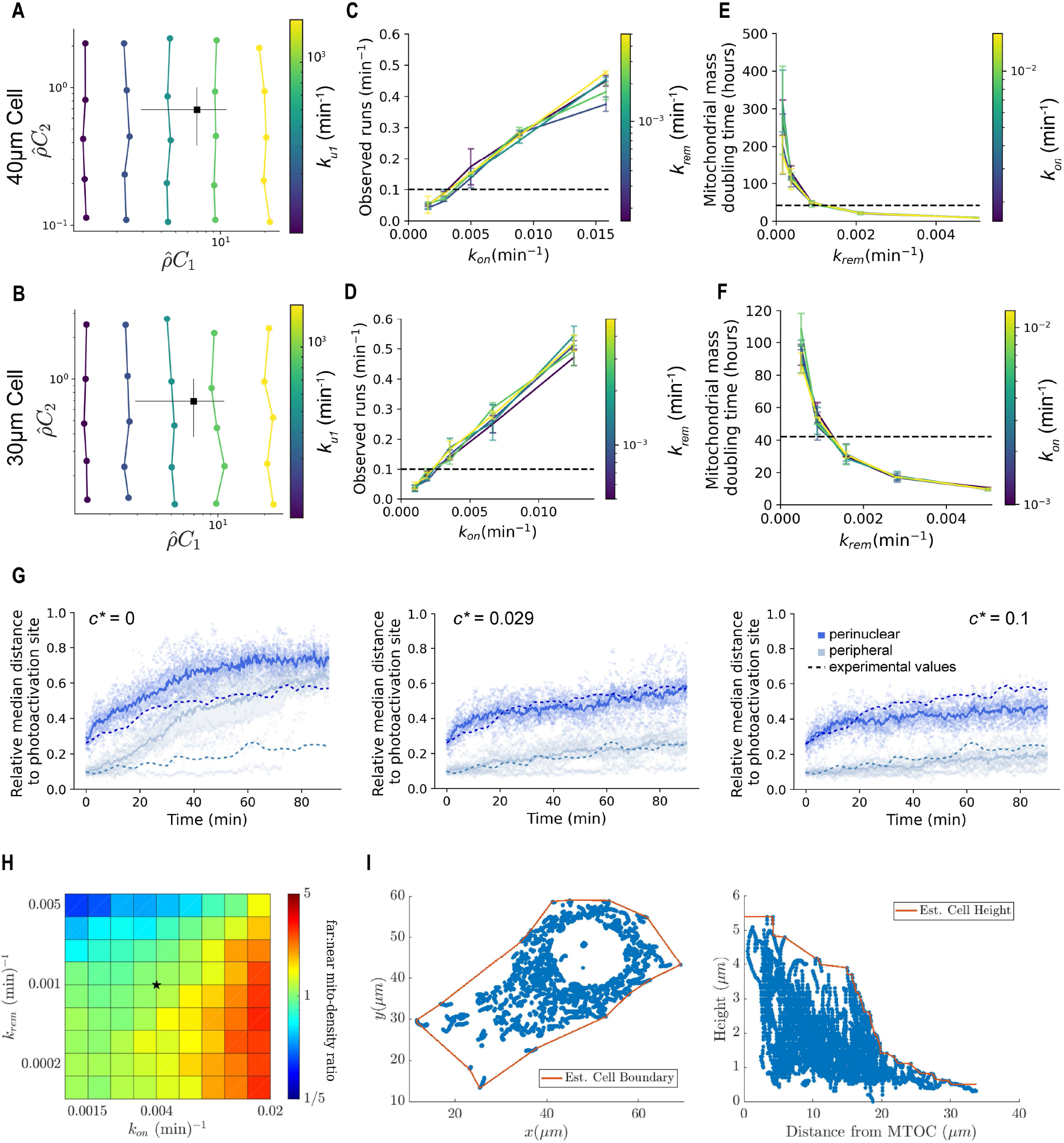
Selection of parameters for the simulation model. Selection of parameters for the simulation model. (A-B). Extraction of fusion parameters for 40 µm (30 µm) cells. Mean-field association constants C1, C2 are calculated for experimental COS-7 networks (obtained using Mitograph) and simulated networks. Colored dots show simulation results for a parameter sweep over several ku,1, ku,2 values. The local tip-tip fusion parameter increases from left to right (colors) while the local tip-side fusion parameter increases bottom to top along each colored line. Black dot shows experimental measurements with standard deviation across different cells indicated by error bars. (C-D). The rate of observed directed runs in simulations is plotted as a function of increasing k_on_ for different values of k_rem_ (colors) in 40 µm (30 µm) cells. The experimentally relevant rate (≈ 0.1 min-1) is indicated by a dashed line. The corresponding on-rate for individual mitochondrial units is k_on_ ≈ 0.0040 min-1 (0.0024 min-1). (E-F). The time to double the mitochondrial mass in simulations is plotted as a function of the turnover rate k_rem_ for different values of the on-rate k_on_ (colors) in 40 µm (30 µm) cells. The experimentally relevant cell division time (≈ 42 hr) is indicated by a dashed line. The corresponding removal rate used in simulations is k_rem_ ≈ 0.0010 min-1 (0.0012 min-1). (G) Simulated ratio of median distances from the photoconversion site for mitochondria labeled with photoconverted signal versus all mitochondria. Dark blue: perinuclear photoconversion site, light blue: peripheral photoconversion site. Dashed lines re-plot the experimental averages from Fig. 1O, vertically offset such that the experimental and simulation curves coincide at time zero. Labeled mitochondria are defined as those containing photoconverted material above a threshold value c* (given as a fraction of the maximum signal per mitochondrial unit). Left: threshold of 0, Middle: threshold value that best fits experimental measurements, Right: higher threshold value. (H) Far:near ratio variation with biogenesis/removal rates (k_rem_) and transport rates (k_on_). Star indicates experimental values. (I) Cellular outline extrapolated from volumetric mitochondrial positioning, showcasing the variability in cell height between the perinuclear and peripheral regions.

**Figure S3:**
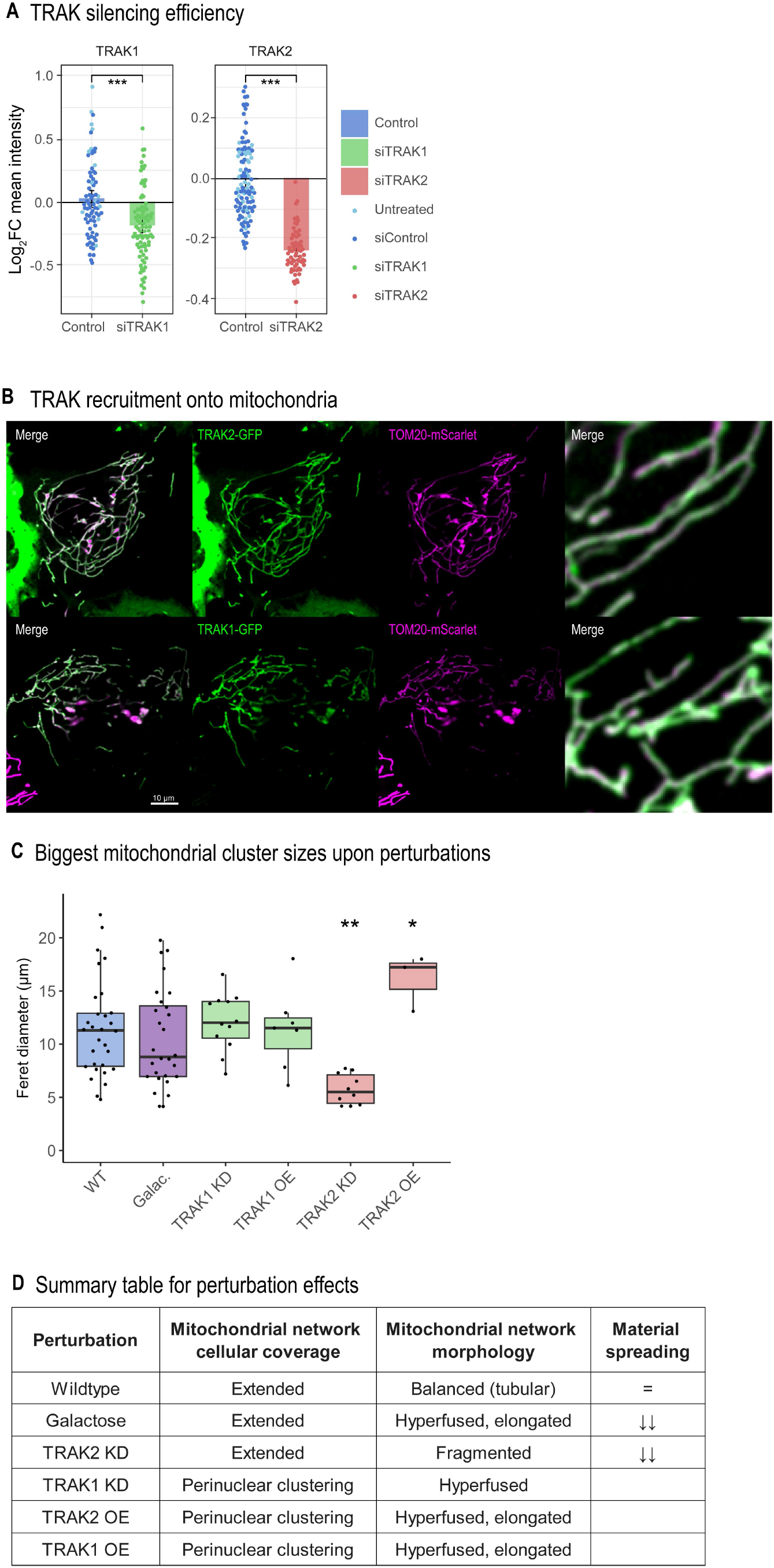
(A) Silencing efficiency of TRAK1 and TRAK2 as quantified by blinded quantification of immunoassayed cells; Ctrl = Control group, including non-treated (dark blue) and control siRNA-treated cells (light blue). P values from a two-sided Wilcoxon rank sum test against controls (p< 0.05*, 0.01**, 0.001***). (B) Representative images of TRAK1 and TRAK2 recruitment into mitochondria upon overexpression, GFP-fusion proteins (green) and mitochondrial TOM20 (magenta). Zoom-in as inset. (C) Feret’s diameter of largest mitochondrial cluster from segmented volumetric networks. P values from a Kruskal-Wallis/Dunn’s test with Bonferroni correction against controls (p< 0.05*, 0.01**, 0.001***). (D) Summary of the main quantified features across perturbations. Material spreading could not be robustly quantified in cells without extended networks due to lack of peripheral mitochondria.

**Movie S1**. Time-lapse widefield fluorescence microscopy of COS-7 cells transfected with a mitochondrial matrix-targeted mEos3.2 construct (48 h). mEos3.2 was photoconverted within regions indicated by dashed circles; green = pre-photoconversion (native), magenta = photoconverted. Cell outline (grey) is derived from the corresponding phase-contrast image. Frames were acquired every 3 min over a total duration of 90 min, displayed at 3 frames/sec. Related to Fig. 1.

**Movie S2**. Time-lapse widefield fluorescence microscopy of COS-7 cells transfected with a mitochondrial matrix-targeted mEos3.2 construct (48 h, green). Cell outline (grey) is derived from the corresponding phase-contrast image. Frames were acquired every 2 sec over a total duration of 15 min, displayed at 20 frames/sec. Arrowheads indicate anterograde (purple) and retrograde (orange) runs. Related to Fig. 1.

